# PiNUI: A Dataset of Protein–Protein Interactions for Machine Learning

**DOI:** 10.1101/2023.12.12.571298

**Authors:** Geoffroy Dubourg-Felonneau, Daniel Mitiku Wesego, Eyal Akiva, Ranjani Varadan

## Abstract

We introduce a novel dataset named **PiNUI**: **P**rotein **I**nteractions with **N**early **U**niform **I**mbalance. PiNUI is a dataset of Protein–Protein Interactions (PPI) specifically designed for Machine Learning (ML) applications that offer a higher degree of representativeness of real-world PPI tasks compared to existing ML-ready PPI datasets. We achieve such by increasing the data size and quality, and minimizing the sampling bias of negative interactions. We demonstrate that models trained on PiNUI almost always outperform those trained on conventional PPI datasets when evaluated on various general PPI tasks using external test sets. PiNUI is available [here](https://linktr.ee/geoffroy.shiru).

## 1 Introduction

Proteins are nanomachines that carry out the majority of cellular processes. They frequently work in concert to carry out their functions. Thus, identifying which proteins interact with one another is key to understanding fundamental cellular processes. Since experimental validation of PPIs is a cumbersome task, prediction typically provides the first step in PPI elucidation. Accurate PPI prediction can help identify disease mechanisms. For example, finding proteins of pathogenic bacteria that interact with human proteins can help in developing drugs that inhibit these interactions.

We recognized that the quality of the training set is paramount when it comes to Protein-Protein Interaction prediction. Factors such as the data power, the strength of supporting evidence for positive labels, and particularly the methodology employed to generate negative labels are all expected to significantly influence the subsequent models’ ability to 1) perform effectively and 2) generalize to real PPI scenarios across different contexts and organisms.

For this reason, we assert that the pursuit of PPI research through Machine Learning necessitates a novel dataset—one characterized by increased sample size and reduced selection bias in negative labels.

In this study, we conduct a comparative analysis between models trained on two widely employed datasets in Machine Learning research for PPI and models trained on PiNUI. This comparison encompasses evaluations on their respective test sets, as well as the assessment on a collection of newly curated test sets from diverse sources, specifically tailored to evaluate cross-species interactions. This extension is a departure from the models’ initial training, which focused solely on intra-species interactions.

## 2 Previous work

### 2.1 Baseline Datasets

We utilize two prominent baseline datasets in this study: Guo’s dataset [1], which focuses on Protein-Protein Interactions (PPI) in yeast, and Pan’s dataset, which centers on PPI in humans. These datasets have been widely adopted in machine learning applications for PPI and serve as reference benchmarks for protein language models, such as PEER [2].

Guo’s dataset is employed for training, validation, and testing, using the predefined splits provided by PEER. Positive interactions are extracted from the Database of Interacting Proteins (DIP). To create negative interactions, random pairs of proteins are selected, ensuring that they belong to different cellular compartments.

Similarly, Pan’s dataset [3] is used for training, validation, and testing, following the splits provided by PEER. Positive interactions are sourced from the Human Protein Reference Database (HPRD). Negative interactions are generated by selecting random pairs of proteins from distinct cellular compartments.

Furthermore, we merged the Guo and Pan datasets to create a composite dataset referred to as Guo+Pan. This combination serves a dual purpose: first, to investigate whether models trained on two intra-species interaction datasets exhibit improved performance in inter-species interactions, and second, to augment the overall dataset size.

## 3 PiNUI

**Motivation**: Our goal is to encourage models to learn PPI prediction primarily from the co-occurrence of input proteins within a pair, rather than relying on the occurrence of individual proteins.

Positive interactions within PiNUI are exclusively sourced from the European Bioinformatics Institute’s intAct [4] interactome, parsed according to the miXML2.5 [5] specification. Initially, our focus is on two organisms: *Homo sapiens* (Taxonomy id: 9606), referred to as Human, and *Saccharomyces cerevisiae* (Taxonomy id: 559292), referred to as Yeast.

### 3.1 Negative set

In both the baseline and PiNUI datasets, the positive set is formed based on the presence of experimental evidence confirming an interaction between pairs. However, there are notable differences in the curation strategy employed for the negative set, which we elucidate as follows:

- We select sequences exclusively from the positive sequence pairs. We do not include sequences from Negatomes or choose random sequences from the organism that are not part of at least one positive pair. This approach is intentional to prevent any negative bias toward sequences that exclusively appear in the negative set.
- In previous studies, we noticed a recurring pattern where a specific sequence frequently appeared in positive pairs while never being part of a negative pair. Conversely, the reverse situation also occurred frequently. This intrinsic bias stemming from the dataset’s structure has the potential to mislead the model into forming an inaccurate association between a particular sequence and a seemingly spurious label frequency. This, in turn, hinders the model from effectively relying on the co-occurrence of sequences as expected. While creating PiNUI, our aim was to maintain a uniform ratio of positive to negative labels per sequence. This means that sequences linked to positive interactions should exhibit a proportional presence in negative interactions. Let *S* be the set of sequences involved in positive interactions. Let *P* = {(*a, b*)| *a, b* ∈ *S*} be the set of positive pairs. In order to guarantee this proportionality, for each positive pair (*a, b*) ∈ *P*, we create one negative pair (*a, x*) where *x* ∈ *S \* {*t*|(*a, t*) ∈ *P, t* ∈ *S*}, and one negative pair (*y, b*) where *y* ∈ *S \* {*t*|(*t, b*) ∈ *P, t* ∈ *S*. While this strategy does introduce label imbalance, it does so systematically on a per-protein basis. As a result, it yields a nearly uniform distribution of (im)balanced labels across proteins, a feature we anticipate will be advantageous for the model.
- We do not omit interologs [6], i.e. PPIs where the two protein partners are homologous to another pair of interaction members, through the process of sequence clustering. The model can learn from such cases, since even small variations in sequence can have a major impact on the structure, and thereby PPI interaction.
- We do not use subcellular localization as a gold standard for negatives. Most of the existing datasets [1][7] use cellular localization annotations or predictions to find negative pairs, assuming that if proteins live in different cell compartments they cannot interact. We argue that while this may be a reasonable constraint in vivo, the identity of the native physical compartment should not preclude interactions between proteins in other contexts such as therapeutics and food systems. We want our dataset to be as general as possible. Proteins in different subcellular localization, thus are presumably not interacting *in vivo*, but could interact in a cell-free environment, or in a different organism (for instance, in case they are heterlogously expressed in a host organism, where localization is not preserved). Furthermore, it is important to note that researchers have demonstrated [8][9][10] the high accuracy of Protein Language Model (LM) in predicting subcellular localization. Systematically labeling pairs of proteins from different subcellular localizations as negatives may inadvertently lead models to learn a proxy for subcellular localization rather than genuine protein interactions. Hence, we advocate for a more inclusive approach in defining negative interactions in order to foster a dataset that better reflects the complexity and versatility of protein-protein interactions.

### 3.2 PiNUI:yeast

To construct PiNUI:yeast we curated all positive and negative interactions provided in the interactome from intAct. There are 43, 966 unique positive interactions and 4 evidence-backed negative interactions that contain proteins of length shorter than 1, 022 (intrinsic limit of the esm representtion model). From this set we have removed all the interactions involving at least one of the test 393 sequences from the test split of Guo provided by PEER, so that we may be able to use this test across multiple methods. Finaly, we generate the negative set following the method described in 3.1.

### 3.3 PiNUI:human

Following the same process as in PiNUI:yeast, the curation of the human interactome left us with 229, 135 positive and 888 negative pairs (before generating the negative set as described in 3.1).

### 3.4 Test sets

- **Human receptors**: For this test set (*N* = 60), a compilation of literature-documented receptors from the human proteome was constructed by integrating a non-redundant dataset of proteins from CellTalkDB [11] and the human gpDB [12]. This compilation was used as a positive set. The negative set was constructed similarily to the Human–non-human test sets, where proteins from each side of the positive pair was replaced by a random counterpart of similar or same organism.
- **Human–non-human**: For testing purposes, we designed and curated two datasets focusing on human–non-human protein interactions. The first dataset (*N* = 390) was constructed through a query of the Research Collaboratory for Structural Bioinformatics’ Protein Data Bank (PDB). We specifically sought experimentally validated structures of heterodimers, where one protein belongs to Human, and the other originates from a non-human organism. Experimental methods were restricted to nuclear magnetic resonance (NMR), Electron-Microscopy (EM) or X-ray diffraction. These instances constituted our positive pairs. To create a negative set, we took all the positive pairs and systematically substituted the human proteins with randomly sampled human proteins. Similarly, we generated another negative set by replacing non-human proteins with randomly chosen non-human proteins, sourced from a pool of bacteria and plant proteins, as these align with the non-human proteins present in the positive set. Our objective here was to demonstrate the model’s ability to predict PPIs where one protein within a pair (the Human one) has representation in the training set, while the other originates from a distinct species. Creating a negative set in such a manner allows us to test that proteins seen in positive pairs can be predicted in a negative pair when the other participant doesn’t interact with them. In other words, we want to test that the positive labels are not solely associated with proteins in positive pairs, but with a combination of interacting proteins. This method yields a 1:2 (positive:negative) label balance; for this reason, we use the AUC-ROC as our main performance metric, as it is still relevant to use on lightly imbalanced datasets as such. The second human-non-human test (*N* = 292) set was generated using a similar approach with orthogonal experimental validations. In this case, we utilized the European Bioinformatics Institute’s intAct database to assemble positive pairs. Our criterion for inclusion was the occurrence of heterodimers, where one protein belongs to Human while the other does not. Notably, the key distinction lays in the difference of experimental methods used to gather PPI evidence, including antitag co-immunoprecipitation, multiple variants of two-hybrid assays, and other methodologies that did not overlap with the PDB dataset.
- **Human-SARS-CoV2**: Similar to Human–non-human, sourced from intact, and specifically restricting non-human proteins to come from SARS-CoV2 (*N* = 1, 492).
- **Human–Yeast**: Similar to Human–non-human, sourced from intact, and specifically restricting non-human proteins to come from Yeast (*N* = 468).
- **Human–Mouse**: Similar to Human–non-human, sourced from RCSB, and specifically restricting non-human proteins to come from Mus musculus (*N* = 78).

## 4 Comparison

In order to demonstrate the benefits of our dataset, we trained the same Neural Network (3-layers MLP) on Guo’s, Pan’s, and PiNUI’s datasets. To represent the protein sequences, we use Meta’s Evolutionary Scale Model [13]. The representations are then concatenated to form the input vector.

In Table 1, a conspicuous trend emerges as models trained on the PiNUI variants outperform their counterparts trained on PEER’s datasets by a significant margin. Notably, the Guo-trained model exhibits commendable performance (.67AUC) on the Guo test set, while the Pan-trained model excels (.93AUC) on the Pan test set. However, these specialized strengths come at the cost of diminished performance on other test sets. In contrast, the PiNUI-trained models, although displaying lower test performance on their respective test sets (.80, .76, .75AUC) compared to the baseline, offer a more representative indication of their overall performance across diverse test sets.

**Table 1:** AUC-ROC scores of 3 baseline models (trained on Guo’s and Pan’s datasets), and 3 equivalent models trained on PiNUI datasets, tested on their 6 respective test sets, and 6 additional test sets from various sources. The value reported is the mean AUC-ROC of all the models trained in a 5-fold cross-validation, and the standard deviation in parenthesis.

|  | Baseline |  |  | PiNUI |  |  |
| --- | --- | --- | --- | --- | --- | --- |
|  | Guo (Yeast) | Pan (Human) | Guo+Pan | PiNUI:yeast | PiNUI:human | PiNUI:yeast+human |
| Guo (Yeast) | 0.668 (0.03) | 0.563 (0.01) | 0.562 (0.03) | 0.568 (0.02) | 0.615 (0.01) | 0.570 (0.01) |
| Pan (Human) | 0.504 (0.04) | 0.930 (0.00) | 0.920 (0.01) | 0.530 (0.03) | 0.710 (0.01) | 0.689 (0.02) |
| Guo+Pan | 0.547 (0.04) | 0.611 (0.01) | 0.632 (0.01) | 0.629 (0.01) | 0.478 (0.00) | 0.595 (0.01) |
| PiNUI:yeast | 0.580 (0.02) | 0.557 (0.00) | 0.543 (0.03) | 0.801 (0.00) | 0.622 (0.00) | 0.753 (0.00) |
| PiNUI:human | 0.546 (0.00) | 0.535 (0.01) | 0.536 (0.00) | 0.523 (0.01) | 0.757 (0.00) | 0.751 (0.00) |
| PiNUI:yeast+human | 0.550 (0.00) | 0.534 (0.00) | 0.533 (0.00) | 0.559 (0.01) | 0.718 (0.00) | 0.751 (0.00) |
| Human receptors | 0.521 (0.02) | 0.534 (0.01) | 0.557 (0.03) | 0.655 (0.04) | 0.732 (0.05) | 0.721 (0.03) |
| Human–non-human1 | 0.500 (0.00) | 0.503 (0.01) | 0.495 (0.00) | 0.533 (0.01) | 0.617 (0.01) | 0.619 (0.00) |
| Human–non-human2 | 0.671 (0.05) | 0.508 (0.01) | 0.509 (0.01) | 0.540 (0.02) | 0.829 (0.02) | 0.838 (0.01) |
| Human-SARS-CoV2 | 0.500 (0.03) | 0.439 (0.01) | 0.433 (0.02) | 0.667 (0.03) | 0.592 (0.01) | 0.609 (0.01) |
| Human–Yeast | 0.561 (0.05) | 0.382 (0.02) | 0.380 (0.03) | 0.746 (0.03) | 0.518 (0.02) | 0.499 (0.01) |
| Human–mouse | 0.820 (0.04) | 0.535 (0.02) | 0.608 (0.10) | 0.612 (0.11) | 0.854 (0.01) | 0.844 (0.03) |

This highlights the importance of the training set’s quality, with a specific emphasis on the methodology employed to generate unobserved negative protein-protein interactions for training purposes. Conversely for experimental biologists looking to leverage ML-models, the study highlights the importance of mitigating unintended bias when designing studies aimed at collection of large datasets for predictive ML-models.

## 5 Conclusion

In conclusion, our analysis demonstrates that models trained on the PiNUI dataset almost in every instance, outperform their counterparts trained on PEER’s datasets when evaluated across a diverse range of test sets. This suggests that the different design approach used to build PiNUI does encourage models to generalize better across various testing scenarios.

The PiNUI dataset is intentionally designed to incorporate a nearly uniform imbalance in its label distribution, challenging models to learn meaningful protein associations primarily from the co-occurrence of input proteins within a pair. This design choice fosters a robust learning process, enabling models to focus on, we hope, genuine protein interactions rather than relying on the mere occurrence of individual proteins.

## References

[1] Yanzhi Guo, Lezheng Yu, Zhining Wen, and Menglong Li. Using support vector machine combined with auto covariance to predict protein–protein interactions from protein sequences. Nucleic Acids Research, 36(9):3025–3030, April 2008.

[2] Minghao Xu, Zuobai Zhang, Jiarui Lu, Zhaocheng Zhu, Yangtian Zhang, Ma Chang, Runcheng Liu, and Jian Tang. Peer: a comprehensive and multi-task benchmark for protein sequence understanding. Advances in Neural Information Processing Systems, 35:35156–35173, 2022.

[3] Xiao-Yong Pan, Ya-Nan Zhang, and Hong-Bin Shen. Large-scale prediction of human proteinprotein interactions from amino acid sequence based on latent topic features. Journal of Proteome Research, 9(10):4992–5001, August 2010.

[4] Noemi del Toro, Anjali Shrivastava, Eliot Ragueneau, Birgit Meldal, Colin Combe, Elisabet Barrera, Livia Perfetto, Karyn How, Prashansa Ratan, Gautam Shirodkar, Odilia Lu, Bálint Mészáros, Xavier Watkins, Sangya Pundir, Luana Licata, Marta Iannuccelli, Matteo Pellegrini, Maria Jesus Martin, Simona Panni, Margaret Duesbury, Sylvain D Vallet, Juri Rappsilber, Sylvie Ricard-Blum, Gianni Cesareni, Lukasz Salwinski, Sandra Orchard, Pablo Porras, Kalpana Panneerselvam, and Henning Hermjakob. The IntAct database: efficient access to fine-grained molecular interaction data. Nucleic Acids Research, 50(D1):D648–D653, November 2021.

[5] Samuel Kerrien, Sandra Orchard, Luisa Montecchi-Palazzi, Bruno Aranda, Antony F Quinn, Nisha Vinod, Gary D Bader, Ioannis Xenarios, Jérôme Wojcik, David Sherman, Mike Tyers, John J Salama, Susan Moore, Arnaud Ceol, Andrew Chatr-aryamontri, Matthias Oesterheld, Volker Stümpflen, Lukasz Salwinski, Jason Nerothin, Ethan Cerami, Michael E Cusick, Marc Vidal, Michael Gilson, John Armstrong, Peter Woollard, Christopher Hogue, David Eisenberg, Gianni Cesareni, Rolf Apweiler, and Henning Hermjakob. Broadening the horizon – level 2.5 of the HUPO-PSI format for molecular interactions. BMC Biology, 5(1), October 2007.

[6] Sven Mika and Burkhard Rost. Protein–protein interactions more conserved within species than across species. PLoS Computational Biology, 2(7):e79, July 2006.

[7] Zhu-Hong You, Keith C. C. Chan, and Pengwei Hu. Predicting protein-protein interactions from primary protein sequences using a novel multi-scale local feature representation scheme and the random forest. PLOS ONE, 10(5):e0125811, May 2015.

[8] Vineet Thumuluri, José Juan Almagro Armenteros, Alexander Rosenberg Johansen, Henrik Nielsen, and Ole Winther. DeepLoc 2.0: multi-label subcellular localization prediction using protein language models. Nucleic Acids Research, 50(W1):W228–W234, April 2022.

[9] Gurpreet Singh, Ravi Tyagi, Anjana Singh, Shruti Kapil, Pratap Kumar Parida, Maria Scarcelli, Dan Dumitru, Nanda Kumar Sathiyamoorthy, Sanjay Phogat, and Ahmed Essaghir. Protein language model for prediction of subcellular localization of protein sequences from gramnegative bacteria (ProtLM.SCL). December 2022.

[10] Geoffroy Dubourg-Felonneau, Arash Abbasi, Eyal Akiva, and Lawrence Lee. Improving protein subcellular localization prediction with structural prediction & graph neural networks. In NeurIPS 2022 Workshop on Learning Meaningful Representations of Life, 2022.

[11] Xin Shao, Jie Liao, Chengyu Li, Xiaoyan Lu, Junyun Cheng, and Xiaohui Fan. CellTalkDB: a manually curated database of ligand–receptor interactions in humans and mice. Briefings in Bioinformatics, 22(4), November 2020.

[12] V. P. Satagopam, M. C. Theodoropoulou, C. K. Stampolakis, G. A. Pavlopoulos, N. C. Papandreou, P. G. Bagos, R. Schneider, and S. J. Hamodrakas. GPCRs, g-proteins, effectors and their interactions: human-gpDB, a database employing visualization tools and data integration techniques. Database, 2010(0):baq019–baq019, August 2010.

[13] Alexander Rives, Joshua Meier, Tom Sercu, Siddharth Goyal, Zeming Lin, Jason Liu, Demi Guo, Myle Ott, C. Lawrence Zitnick, Jerry Ma, and Rob Fergus. Biological structure and function emerge from scaling unsupervised learning to 250 million protein sequences. Proceedings of the National Academy of Sciences, 118(15):e2016239118, 2021.

